# COVID-19-related research data availability and quality according to the FAIR principles: A meta-research study

**DOI:** 10.1101/2023.11.14.566998

**Authors:** Ahmad Sofi-Mahmudi, Eero Raittio, Yeganeh Khazaei, Javed Ashraf, Falk Schwendicke, Sergio E. Uribe, David Moher

## Abstract

**Background:** As per the FAIR principles (Findable, Accessible, Interoperable, and Reusable), scientific research data should be findable, accessible, interoperable, and reusable. The COVID-19 pandemic has led to massive research activities and an unprecedented number of topical publications in a short time. There has not been any evaluation to assess if this COVID-19-related research data complied with FAIR principles (or FAIRness) so far.

**Objective:** Our objective was to investigate the availability of open data in COVID-19-related research and to assess compliance with FAIRness.

**Methods:** We conducted a comprehensive search and retrieved all open-access articles related to COVID-19 from journals indexed in PubMed, available in the Europe PubMed Central database, published from January 2020 through June 2023, using the *metareadr* package. Using *rtransparent*, a validated automated tool, we identified articles that included a link to their raw data hosted in a public repository. We then screened the link and included those repositories which included data specifically for their pertaining paper. Subsequently, we automatically assessed the adherence of the repositories to the FAIR principles using FAIRsFAIR Research Data Object Assessment Service (F-UJI) and *rfuji* package. The FAIR scores ranged from 1–22 and had four components. We reported descriptive analysis for each article type, journal category and repository. We used linear regression models to find the most influential factors on the FAIRness of data.

**Results:** 5,700 URLs were included in the final analysis, sharing their data in a general-purpose repository. The mean (standard deviation, SD) level of compliance with FAIR metrics was 9.4 (4.88). The percentages of moderate or advanced compliance were as follows: Findability: 100.0%, Accessibility: 21.5%, Interoperability: 46.7%, and Reusability: 61.3%. The overall and component-wise monthly trends were consistent over the follow-up. Reviews (9.80, SD=5.06, n=160), and articles in dental journals (13.67, SD=3.51, n=3) and Harvard Dataverse (15.79, SD=3.65, n=244) had the highest mean FAIRness scores, whereas letters (7.83, SD=4.30, n=55), articles in neuroscience journals (8.16, SD=3.73, n=63), and those deposited in GitHub (4.50, SD=0.13, n=2,152) showed the lowest scores. Regression models showed that the most influential factor on FAIRness scores was the repository (R^2^=0.809).

**Conclusion:** This paper underscored the potential for improvement across all facets of FAIR principles, with a specific emphasis on enhancing Interoperability and Reusability in the data shared within general repositories during the COVID-19 pandemic.

## Introduction

The COVID-19 pandemic introduced a significant shift in the scientific publishing ecosystem, catalyzed by the urgency of sharing findings in a rapidly evolving global health crisis (1). This led to an unprecedented proliferation of preprint publications and open-access materials, allowing researchers worldwide to access both peer-reviewed and non-peer-reviewed findings freely (2). Open-access publications are just one aspect of a larger, comprehensive movement: open science (3). Some funders and journals, such as CIHR (4), NIH (5), BMJ (6), and PLOS (7), have aligned themselves with open science. Central to open science are three components: open protocols, open-access publications, and open data; collectively, they enhance transparency, collaboration and dissemination (8).

Data openness is the cornerstone of research validation and replication, fortifying scientific credibility (9). Precise, exhaustive datasets form the bedrock on which scientific conclusions rest and inform the development of further research. In contrast, a major issue during the COVID-19 pandemic was the lack of high-quality, timely, and reliable data, partially feeding the burgeoning “infodemic” (10), where an excessive amount of information, including false or misleading content, is circulated in digital and physical spaces. Inaccurate or insufficient data can lead to skepticism and mistrust toward research findings, eroding public confidence and impeding a science-informed response (11).

To optimally utilize open research data, it must align with the FAIR principles, i.e., that data are Findable, Accessible, Interoperable, and Reusable (12). These criteria foster better data utility, extending its applicability beyond the original work and facilitating the exploration of different theories, substantiation of claims, probing of debates, prevention of unnecessary duplication, and deriving fresh knowledge from existing data (13). While privacy concerns may impede complete data openness, sharing metadata can be a partial but meaningful substitute (14). Metadata can provide insights into the nature of the data and its structure, facilitating interpretation and usability (15).

Notably, recent studies demonstrate that data sharing as the first requirement for open data remains sparse in medical research (16), and that shared data often fail to meet the FAIR principles (17). For assessing research integrity, better decision-making, gaining public trust, and future preparedness, it is essential to clarify the data quality generated throughout the COVID-19 pandemic (18). Thus, this study aimed to assess the adherence of COVID-19-related research data to the FAIR principles (or FAIRness), a critical step towards improving data quality and trust in scientific outputs.

## Methods

The protocol of this study was deposited prior to beginning the study on the Open Science Framework (OSF) website (https://doi.org/10.17605/OSF.IO/XAYP9). All the codes and the data related to the study have been shared via its OSF repository (https://doi.org/10.17605/OSF.IO/YMD6W) and GitHub (https://github.com/choxos/covid-fairness) at the time of submission of the manuscript. Deviations from the protocol are available in Appendix 1.

### Data Sources and Study Selection

First, we searched for all open access PubMed-indexed records available in the Europe PubMed Central (EPMC) database from January 1, 2020 (when Chinese authorities announced the new virus), to June 30, 2023, using the *europepmc* package (19) in R (20). EPMC includes all records available through PubMed and PubMed Central, and allows the retrieval of records automatically. Since the automated tools we used were optimized for the English language, only open-access English papers were included. We used the following query to identify all open-access PubMed-indexed papers in English from the beginning of 2020 until the end of June 2023.

To identify COVID-19-related articles, we selected articles with PubMed ID (PMID) in the LitCovid database (https://ncbi.nlm.nih.gov/research/coronavirus). LitCovid, sponsored by the National Library of Medicine, is a curated literature hub to track up-to-date COVID-19-related scientific information in PubMed. LitCovid is updated daily with newly identified relevant articles organized into curated categories. It uses machine learning and deep learning algorithms (21–23). As we were interested in subgroup analyses along study types, we further used EPMC’s *pubType* column to detect reviews (“review|systematic review|meta-analysis|review-article”), research articles (“research-article”) and letters (“letter”). Since EPMC’s categorization for randomized trials was deemed not to be sensitive enough (24–26) and did not provide any category for all observational studies, we used the L·OVE (Living OVerview of Evidence, https://iloveevidence.com) database to detect randomized trials and observational studies and classify them as such. L·OVE, powered by Epistemonikos Foundation, is an open platform that maps and organizes the best evidence in various medical and health sciences fields (27). We applied the “Reporting data” filter on the L·OVE website to detect PMIDs of RCT studies in our dataset. Then, we downloaded all identified open-access COVID-19-related available records in XML full-text format using the *metareadr* package (28) from the EPMC database.

### Data Extraction

We used the *rtransparent* package (29) for programmatically assessing data availability in the included studies. The reliability of this package has previously been validated with an accuracy of 94.2% (89.7%–97.9%) in detecting the data availability of assessed papers (30). The *rtransparent* uses the *oddpub* package (31) for detecting data-sharing statements in XML files of papers. Briefly, the *oddpub* package uses regular expressions to identify whether an article mentions a) a general database in which data are frequently deposited (e.g., figshare); b) a field-specific database in which data are frequently deposited (e.g., dbSNP); c) online repositories in which data/code are frequently deposited (e.g., GitHub); d) language referring to commonly shared file formats (e.g., csv); e) language referring to the availability of data as a supplement (e.g., “supplementary data”); and f) language referring to the presence of a data sharing statement (e.g., “data availability statement”). It finally checks whether these were mentioned in the context of positive statements (e.g., “can be downloaded”) or negative statements (e.g., “not deposited”) to produce its final adjudication. This adjudication indicates whether a data sharing statement is present, which aspect of data sharing was detected (e.g., mention of a public database), and then extracts the phrase in which this was detected.

Our previous study showed low FAIRness of data provided in field-specific databases and supplements (17). This is due to a lack of some properties that reduce FAIRness, such as the lack of an identifier to the dataset, non-machine-readable metadata, and the use of non-general file formats in field-specific databases. Therefore, we focused on studies that provided a link to a public database for their data. Another reason was to reduce the burden of work that added little to our study and helped automatize the workload.

After filtering the studies that provided their data in a general-purpose repository (limited to the ones that were defined and detected by the *oddpub* package, the list of these repositories is available in Appendix 2), we searched for the URL to their dataset in their full-text XML files. To do this, we used keywords related to general-purpose databases and identified every URL that contained one of the keywords. These keywords are available in Appendix 3. After obtaining all the possible URLs to datasets, we manually screened the links. We included a URL only when it belonged to that specific study, i.e., we excluded URLs to general datasets notably, the COVID-19 Data Repository by Johns Hopkins University, the COVID Chest X-Ray dataset by IEEE, and Covid-19 Data in the United States by New York Times. The most relevant and functioning link was selected based on automatically extracted data availability statements, and full-text was consulted if necessary for the decision. Comprehensive details of our approach of including URLs are available in Appendix 2.

We used the Scimago Journal & Country Rank (SJR, https://www.scimagojr.com) to extract the SJR score, H-index, publisher, subject area, and category of the journals.

### FAIRness assessment

FAIR Principles include four main components about how the shared data/metadata should be: Findable, Accessible, Interoperable, and Reusable (12). These four components are divided into 10 subcomponents (F1-F4, A1-A2, I1-I3, and R1) and five sub-subcomponents. To measure the level of FAIRness, a tool named FAIRsFAIR Research Data Object Assessment Service (F-UJI) has been developed by the FAIRsFAIR project (32). F-UJI is a web service to programmatically assess the FAIRness of research data objects based on metrics developed by the FAIRsFAIR project. It checks each component and subcomponents of FAIRness and assigns scores for each metric and an overall score. The lowest score for each component is 0 and the highest ranges from 3–8; the overall score ranges between 1 (because all of them have URLs, FsF-F1-01D=1) and 22. The metrics, scores, and definitions of each metric are illustrated in Appendix 4.

After finalizing the URLs, we used the F-UJI tool to automatically assess the FAIRness of each dataset. This software is based on Python. We used the *rfuji* package (33) in R, which is an R application programming interface (API) client for F-UJI. The workflow of running each software is available in Appendix 5.

### Analysis

We reported the general characteristics of papers that had shared their data in a general-purpose repository. For the FAIRness assessment, we performed a descriptive analysis of compliance with FAIR metrics. FAIR-level differences between different journals and trends over time were explored. We established a categorization system comprising four compliance levels with FAIR principles for each component of FAIR: 0: incomplete; 1: initial; maximum score: advanced; every other score between initial and advanced: moderate.

We performed the Kruskal-Wallis rank sum test to compare the level of FAIRness between different article types, journal subject areas, and repositories. To determine the most influential factor among article type, journal subject area, and repository, we ran different regression models, adjusted for the number of citations and SJR score. Then, we compared the R^2^ of the models. The factor that was in the model with the highest R^2^ was considered the most influential factor.

## Results

### General characteristics

From January 1, 2020 until April 15, 2023, there were 345,332 COVID-19-related articles, including open-access and non-open-access publications. Of those, 257,348 (74.5%) had available full text from the EPMC. However, 7 (<0.01%) of these open-access articles were not downloadable because of technical issues and were excluded from our analyses. Consequently, the sample included 257,341 full-text articles.

Of these, 20,873 (8.1%) were detected to have shared their data; of these, 8,015 (38.4%) had shared their data in a general-purpose repository. After screening the URLs, 6,180 URLs were included.

Out of those articles which shared their data in a general-purpose repository, 746 (12.1%) were published in 2020, 2,394 (38.7%) in 2021, 2,466 (39.9%) in 2022, and 574 (9.3%) in 2023 (censored data) by April 15. More than 9 in 10 of the papers (n=5,580, 90.3%) were research articles, followed by reviews (n=182, 2.9%), observational studies (n=121, 2.0%), and RCTs (n=107, 1.7%), and one percent (n=64) were letters (Figure 1). The papers were from 1,067 different journals, with the top three being PLoS ONE (n=816, 13.2%), Scientific Reports, (n=327, 5.3%), and Nature Communications (n=209, 3.4%).

**Figure 1.**
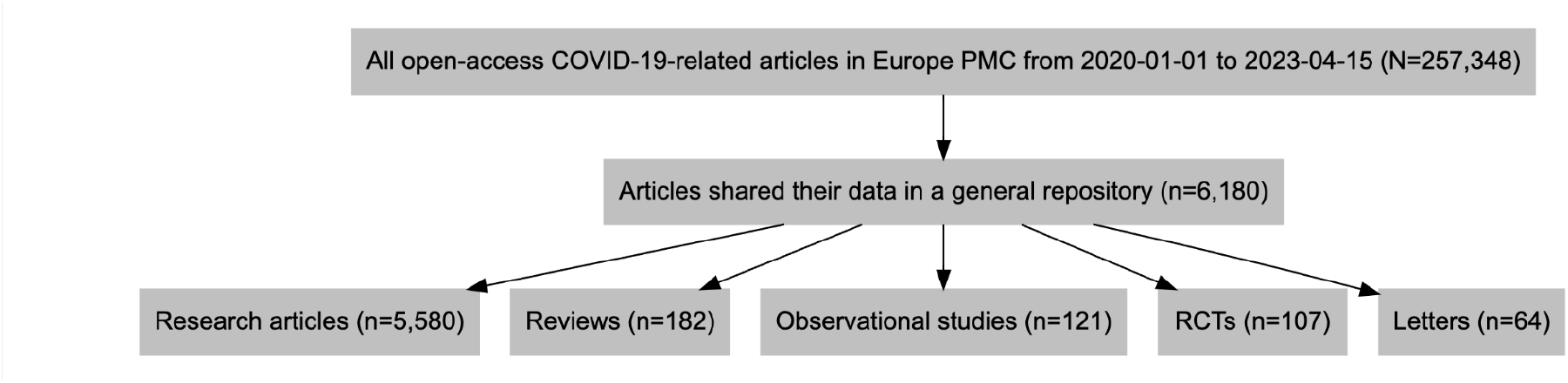
The flow diagram of the study.

### FAIRness results

The FAIRness score for 480 repositories was 1, meaning either the repository was inaccessible or had nothing in it. We eliminated these from our analyses. Therefore, our final analyses were performed on the FAIRness results of 5,700 repositories.

The mean (standard deviation, SD) level of compliance with FAIR metrics was 9.4 (4.88). The mean for each metric was as follows: Findability: 4.3 (1.85) of 7; Accessibility: 1.2 (0.49) of 3; Interoperability: 1.3 (1.30) of 4; and Reusability: 2.6 (1.56) of 8. The compliance by metric is shown in Table 2.

**Table 2.** Summary of FAIR metrics.

| Metric | Mean (SD) | Range | Possible scores |
| --- | --- | --- | --- |
| **FAIR** | 9.4 (4.88) | 1.5–19.0 | 1–22 |
| **<u>F</u>indable** | 4.3 (1.85) | 1.5–7 | 1–7 |
| *F1. (Meta)data are assigned a globally unique and persistent identifier.* | 1.5 (0.50) | 1–2 | 1–2 |
| *F2. Data are described with rich metadata.* | 1.1 (0.66) | 0–2 | 0–2 |
| *F3. Metadata clearly and explicitly include the identifier of the data they describe.* | 0.2 (0.41) | 0–1 | 0–1 |
| *F4. (Meta)data are registered or indexed in a searchable resource.* | 1.4 (0.52) | 0–2 | 0–2 |
| **<u>Accessible</u>** | 1.2 (0.49) | 0–3 | 0–3 |
| *A1. (Meta)data are retrievable by their identifier using a standardized communications protocol.* | 1.2 (0.49) | 0–3 | 0–3 |
| **<u>Interoperable</u>** | 1.3 (1.30) | 0–4 | 0–4 |
| *I1. (Meta)data use a formal, accessible, shared, and broadly applicable language for knowledge representation.* | 1.0 (0.92) | 0–2 | 0–2 |
| *I2. (Meta)data use vocabularies that follow FAIR principles.* | 0.1 (0.24) | 0–1 | 0–1 |
| *I3. (Meta)data include qualified references to other (meta)data.* | 0.3 (0.45) | 0–1 | 0–1 |
| **<u>Reusable</u>** | 2.6 (1.56) | 0–7 | 0–8 |
| *R1. (Meta)data are richly described with a plurality of accurate and relevant attributes.* | 0.7 (0.60) | 0–3 | 0–2 |
| *R1.1. (Meta)data are released with a clear and accessible data usage license.* | 0.8 (0.92) | 0–2 | 0–2 |
| *R1.2. (Meta)data are associated with detailed provenance.* | 1.0 (0.10) | 0–1 | 0–2 |
| *R1.3. (Meta)data meet domain-relevant community standards.* | 0.2 (0.37) | 0–1 | 0–1 |

The percentages of moderate or advanced compliance were as follows: Findability: 100.0%, Accessibility: 21.5%, Interoperability: 46.7%, and Reusability: 61.3% (Figure 2).

**Figure 2.**
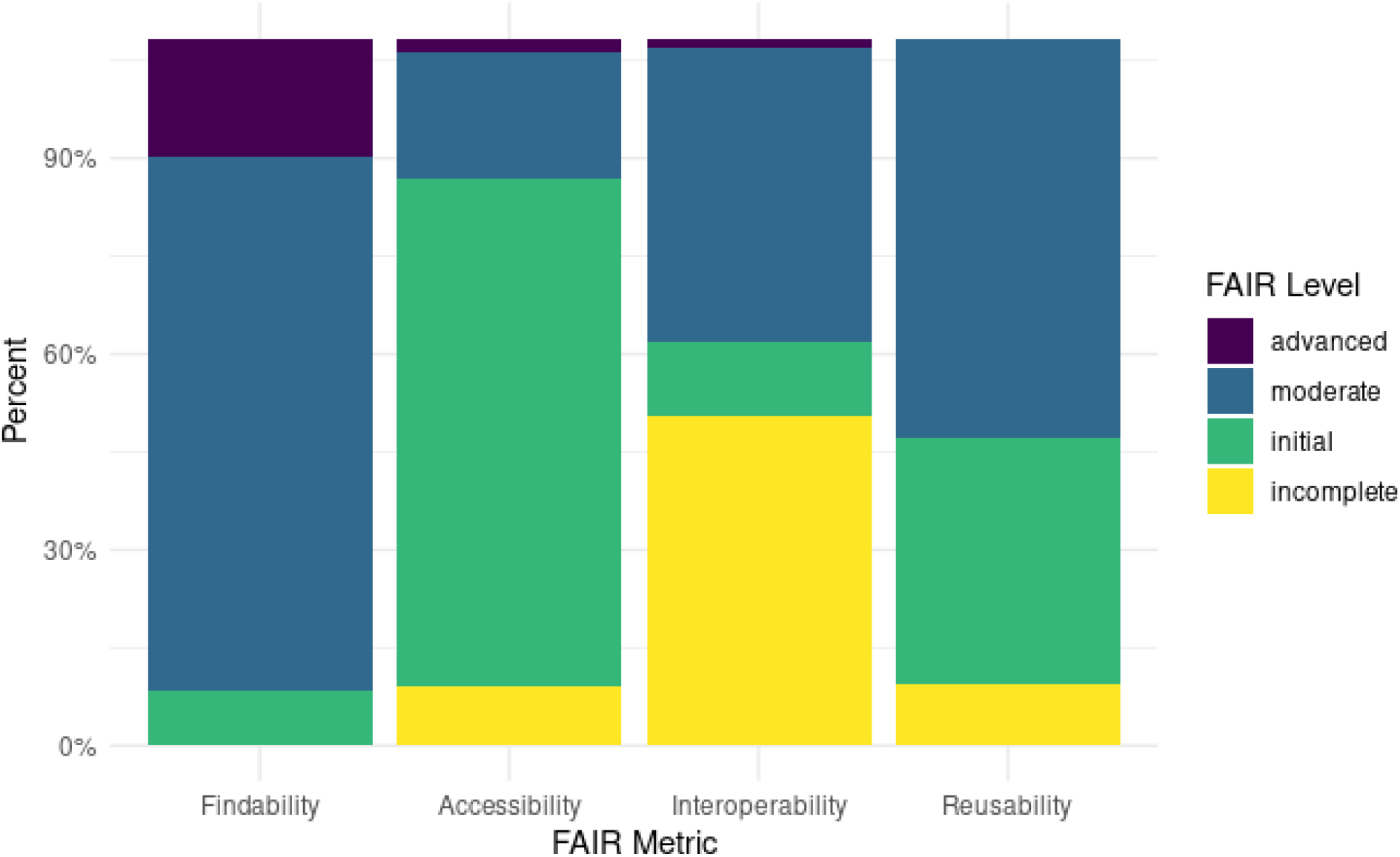
FAIR level by metric (percentage).

Figure 3 shows the yearly mean score in each component of FAIR. All components show decreasing trends. However, monthly trends for the overall and component-wise trends were consistent over the follow-up (Figure 4).

**Figure 3.**
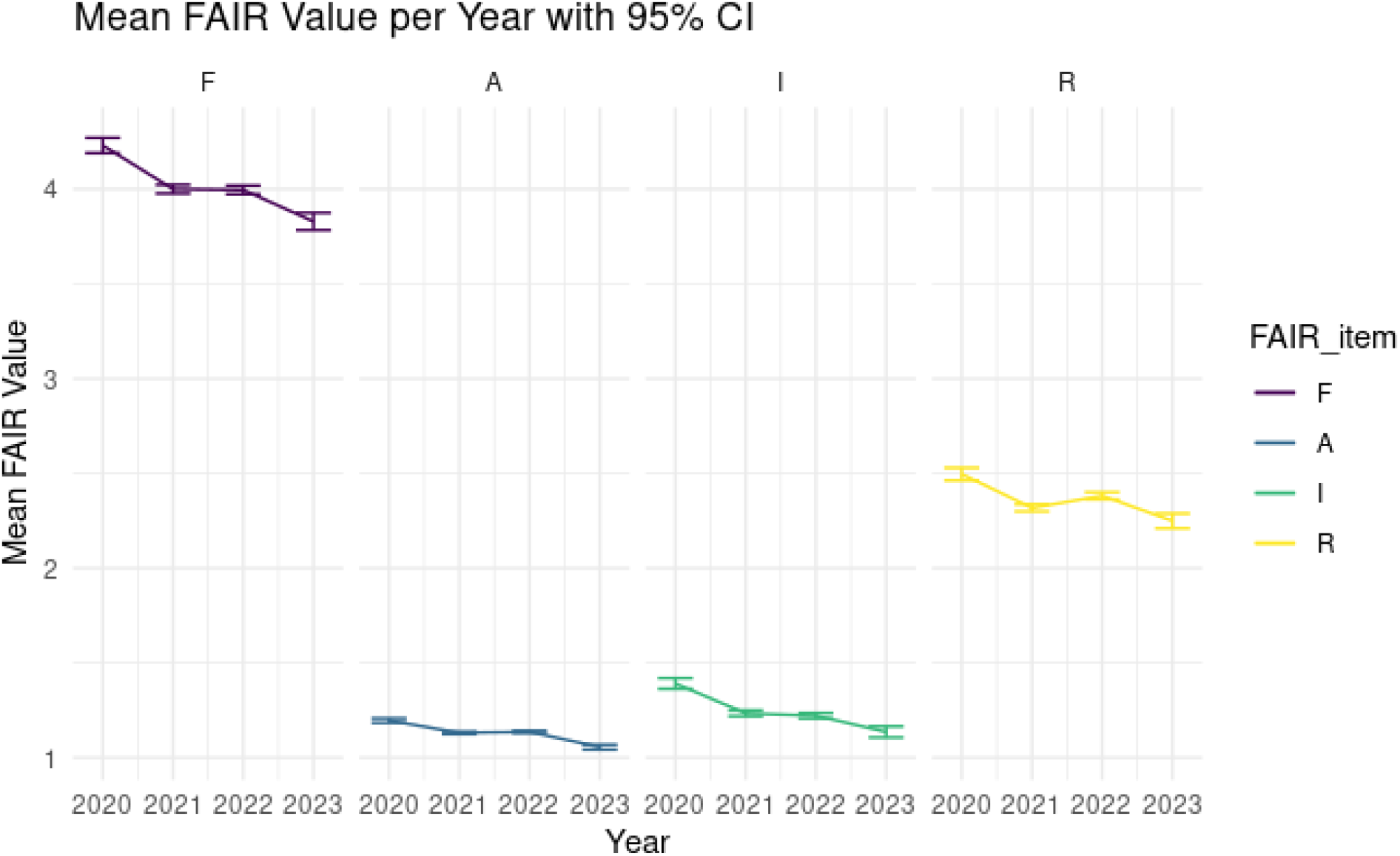
Yearly trends for components of FAIR.

**Figure 4.**
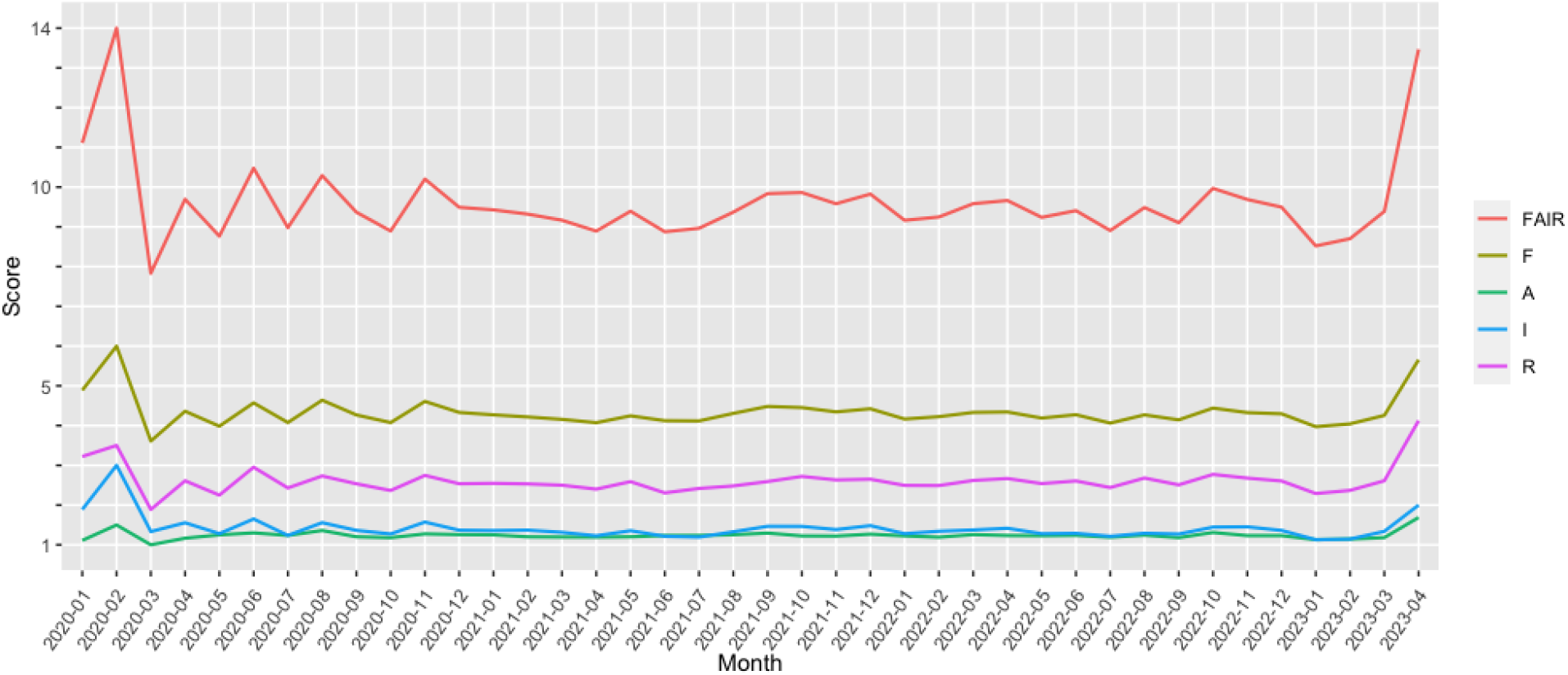
Monthly trend for FAIR score and its components.

### FAIRness by article type

Reviews had the highest mean FAIRness score (9.80, SD=5.06, n=160), whereas research letters had the lowest score (7.83, SD=4.30, n=55). The Kruskal-Wallis rank sum test showed a *P*-value of 0.15 for the difference between the groups. Table 3 shows the detailed information.

**Table 3.** FAIRness score for each article type (mean and SD).

| Article type | Mean FAIRness score (SD) |
| --- | --- |
| Reviews (n=160) | 9.80 (5.06) |
| Observational studies (n=115) | 9.37 (4.87) |
| RCTs (n=90) | 9.04 (4.20) |
| Research letters (n=55) | 7.83 (4.30) |
| Other research articles (n=5,280) | 9.40 (4.89) |
| *P*-value | 0.15 |

### FAIRness by journal subject area

Articles in dental journals had the highest mean FAIRness score (13.67, SD=3.51, n=3), whereas articles in neuroscience journals had the lowest score (8.16, SD=3.73, n=63). The Kruskal-Wallis rank sum test showed a *P*-value of <0.001 for the difference between the groups. Table 4 shows the detailed information.

**Table 4.**
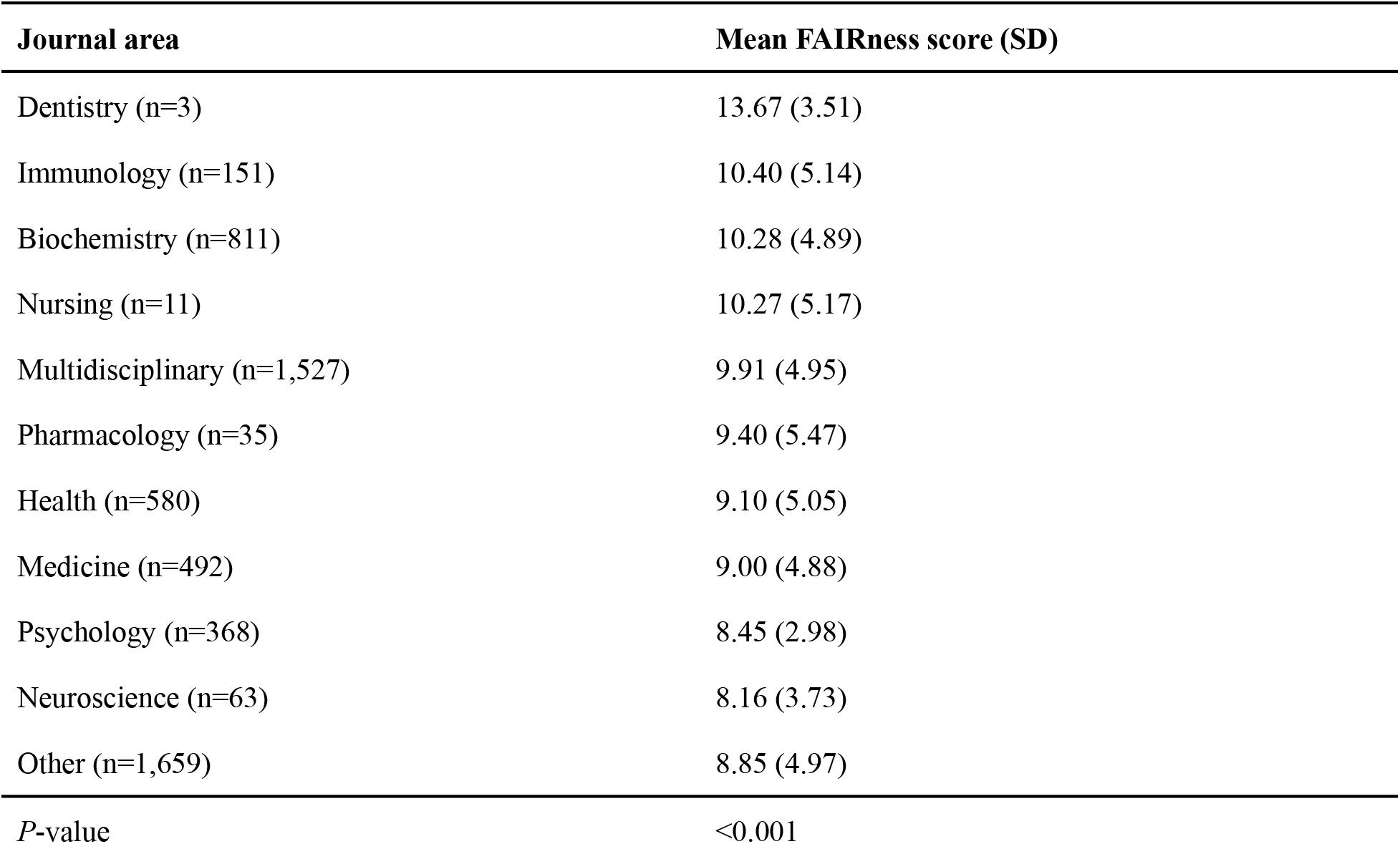
FAIRness score for each journal subject area (mean and SD).

### FAIRness by repository

Harvard Dataverse had the highest mean FAIRness score (15.79, SD=3.65, n=244), whereas GitHub had the lowest score (4.50, SD=0.13, n=2,152). The Kruskal-Wallis rank sum test showed a *P*-value of <0.001 for the difference between the groups. Table 5 shows the detailed information.

**Table 5.** FAIRness score for each repository (mean and SD).

| Repository | Mean FAIRness score (SD) |
| --- | --- |
| Harvard Dataverse (n=244) | 15.79 (3.65) |
| figshare (n=740) | 14.57 (3.50) |
| Mendeley (n=349) | 13.90 (1.17) |
| GigaDB (n=7) | 13.79 (1.95) |
| Zenodo (n=897) | 13.76 (2.28) |
| Dryad (n=140) | 13.73 (4.31) |
| OSF (n=1,134) | 8.43 (2.16) |
| Dataverse NL (n=2) | 7.00 (7.78) |
| GitHub (n=368) | 4.50 (0.13) |
| *P*-value | <0.001 |

### The most influential factor

The R^2^ for the repository was the highest (R^2^=0.809). The R^2^ for the model with all these three factors was 0.812 (Table 6). The *P*-value for the number of citations and SJR score in all models was above 0.29.

**Table 6.**
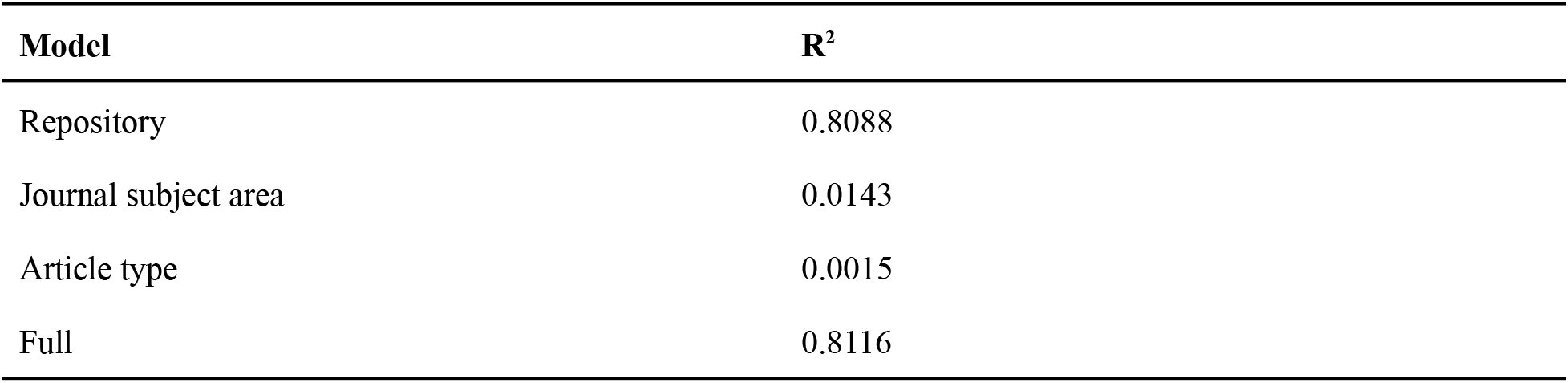
R^2^ for different linear regression models, adjusted for the number of citations and SJR score.

| Model | $R^2$ |
| --- | --- |
| Repository | 0.8088 |
| Journal subject area | 0.0143 |
| Article type | 0.0015 |
| Full | 0.8116 |

## Discussion

Our study aimed to scrutinize FAIR principles adherence in data shared by COVID-19-related articles. We analyzed the data sharing programmatically from 257,348 COVID-19-related articles indexed in PubMed. Out of the articles reviewed, 8.1% were programmatically identified to share their data. Of those, 38.4% utilized a general-purpose repository for data sharing. Finally, we were able to identify functioning links to the general-purpose repositories for 6,180 articles, representing 2.4% of the total number of articles. In those, the overall average FAIR compliance score was 9.4. Compared to the highest possible score in each component, the largest deficiencies were in Reusability, while Findability scored the highest. No considerable changes in the FAIRness compliance were detected over follow-up.

Notable differences in FAIRness compliance emerged based on the repository and journal subject area, but interestingly not based on the type of article. Harvard Dataverse led repository rankings with a mean FAIRness score of 15.8, while GitHub scored the lowest at 4.5. In addition, the regression analysis implied that repository had the greatest impact on the FAIRness of data shared. These differences likely stem from how FAIR principles have been implemented in the structure and workflows within each repository. For instance, in Harvard Dataverse, the principles are implemented systematically to metadata (34), whereas in GitHub, following FAIR principles is much more up to people sharing their data. For instance, data shared via GitHub lacks DOI, and archiving one’s data and materials in general repositories is recommended (35). It is likely that differences in FAIRness by journal subject area are driven by the repository chosen because there were no differences in FAIRness according to the type of article.

As demonstrated through the results of the study, the sharing of data in the medical and healthcare research community remains inappropriately low. This suboptimal adherence to the FAIRness principles has several implications for COVID-19 research. First, it makes it difficult for other researchers to find and use the data. Second, it reduces the reproducibility of research findings. Third, it limits the potential for data reuse and collaboration (12,36,37). Similar studies by (38) and (30) found that only 20% and 18.3% of biomedical articles available in PubMed had a data-sharing statement available in PubMed-based research published between 2015 and 2017 had data available. The absence of transparency in scientific research leads to serious issues with reproducibility, primarily due to the unavailability of data and code. Such opacity significantly hinders a clear understanding of the research methodology and applicability. Considering the public health importance of COVID-19 and the public funding for COVID-19 research, it is disappointing that the essential ingredients necessary for determining the robustness of research publications cannot be validated. Consequently, errors or deficiencies in research design, analysis, reporting, and interpretation persist, even in articles from top-tier journals (39). However, despite the potential benefits of data sharing, its impact on encouraging peer reanalysis has been minimal so far (40).

Despite the privacy constraints inherent in medical and healthcare research, the guiding principle should be to keep data as open as possible yet as closed as necessary (41). Existing guidelines elucidate various advantages, such as enhancing drug safety and efficacy monitoring. This, in turn, spurs research innovation and facilitates secondary analyses for addressing new scientific questions (42). Moreover, adhering to open science practices like data and code sharing in accordance with FAIR principles not only boosts public trust but also fosters greater public engagement in scientific research, data collection, and even research funding (43). Our study’s finding of a low percentage of articles with openly accessible data further underscores the need to restructure incentives for encouraging open scientific practices among researchers. For instance, the Royal Society recommends that “assessment of university research should reward the development of open data on the same scale as journal articles and other publications” (44). Furthermore, the San Francisco Declaration on Research Assessment recommends that funding agencies, institutions, and publishers take into account the significance and influence of all research outputs, encompassing datasets and software, when conducting research evaluations (45). Similarly, the Hong Kong Principles for evaluating researchers endorse the sharing of data and code as an essential component of the publication process (46).

### Strengths and Limitations

To the best of the authors’ knowledge, this is the first study to programmatically assess the application of FAIR principles to COVID-19 research since its official announcement by the Chinese government. Given the recent advancements in algorithmic development, our study offers an initial machine evaluation of the FAIRness of COVID-19 research data. Our study also notes several mentionable strengths: utilizing guidelines like the FAIR principles, we offer a comprehensive evaluation of research data generated in the context of the COVID-19 epidemic. Transparency was maintained throughout—from the idea’s inception to the research submission. The study protocol was pre-published on the OSF website. All relevant codes and data were shared through OSF and GitHub upon manuscript submission. Our approach also incorporates programmatic detection of data availability and repository (30).

A limitation of our study can be attributed to the fact that the study sample focuses solely on articles available from the European PMC database, representing 74.5% of all COVID-19-related medical publications indexed in Pubmed. Moreover, the algorithms from the *rtransparent* package in R were developed before the pandemic, potentially affecting their accuracy in detecting data sharing compared to studies on topics preceding the COVID-19 outbreak. We also only investigated data shared via general repositories, so our findings are directly generalizable to studies sharing data, e.g., as supplementary material on the journal’s website. Finally, one must note that we only evaluated the FAIRness of shared, findable and accessible data, which we were able to identify with our identification strategy. Thus, most research COVID-19-related research data remains unfindable and inaccessible, so that its FAIRness cannot be evaluated systematically.

## Conclusion

Our findings highlight room for improvement in all components of FAIR, but particularly in terms of Interoperability and Reusability, in the data shared in general repositories during the COVID-19 pandemic. Repository chosen had the biggest impact on FAIRness compliance. Enhanced availability of high-quality open research data would bolster confidence in scientific findings and interpretations, thereby narrowing the information gap among researchers, clinicians, and the general populace. Utilizing FAIR principles would thus, for sure, facilitate both human and machine accessibility to research data, thereby augmenting the efficacy of tools designed to navigate intricate, multi-dimensional health-related data. Such machine-actionable data, offering real-time insights, would fortify the emergence of data-driven medicine and ultimately advance healthcare research goals, thereby elevating overall health and quality of life. Joint efforts involving all stakeholders in scientific publishing; researchers, editors, publishers, and funders; accompanied by data repositories are very welcome. Universities also have an important role. This is where the next generation of researchers are currently learning. Incorporating more visible training on open science, including the FAIR principles, is likely be pay dividends to increasing data and code availability and reuse.

## Notes

### Competing Interest Statement

The authors have declared no competing interest.

